# Integrating plant stoichiometry and feeding experiments: state-dependent forage choice and its implications on body mass

**DOI:** 10.1101/2021.02.16.431523

**Authors:** Juliana Balluffi-Fry, Shawn J. Leroux, Yolanda F. Wiersma, Isabella C. Richmond, Travis R. Heckford, Matteo Rizzuto, Joanie L. Kennah, Eric Vander Wal

**Affiliations:** Department of Biology, Memorial University of Newfoundland, St. John’s, Canada; Department of Biological Sciences, University of Alberta, Edmonton, Canada

**Keywords:** herbivory, plant quality, snowshoe hare, foraging ecology, ecological stoichiometry

## Abstract

Intraspecific feeding choices account for a large portion of herbivore foraging in many ecosystems. Plant resource quality is heterogeneously distributed, affected by nutrient availability and growing conditions. Herbivores navigate landscapes, making feeding decisions according to food qualities, but also energetic and nutritional demands. We test three non-exclusive foraging hypotheses using the snowshoe hare (*Lepus americanus*): 1) herbivores feeding choices and body conditions respond to intraspecific plant quality variation, 2) feeding responses are mitigated when energetic demands are high, and 3) feeding responses are inflated when nutritional demands are high. We measured black spruce (*Picea mariana*) nitrogen, phosphorus, and terpene compositions, as indicators of quality, within a snowshoe hare trapping grid and found plant growing conditions to explain spruce quality variation (R^2^ < 0.36). We then offered two qualities of spruce (H1) from the trapping grid to hares in cafeteria-style experiments and measured their feeding and body condition responses (n = 75). We proxied energetic demands (H2) with ambient temperature and coat insulation (% white coat) and nutritional demands (H3) with the spruce quality (nitrogen and phosphorus content) in home ranges. Hares that preferred higher-quality spruce lost less weight during experiments (p = 0.018). The results supported our energetic predictions: hares in colder temperatures and with less-insulative coats (lower % white) consumed more spruce and were less selective towards high-quality spruce. Collectively, we found variation in plant growing conditions within herbivore home ranges substantial enough to affect herbivore body conditions, but any plant-herbivore interactions are also mediated by animal energetic states.

## Introduction

Herbivores have evolved to maximize the energetic and nutritional gain from foraging across heterogenous distributions of plant food (Vivas et al. 1991; Jensen et al. 2012). It is widely accepted that plant species differ in quality (Rodgers and Sinclair 1997), i.e. composition, but even within a species, plant quality changes across landscapes (Leroux et al. 2017). Microclimates, landscape features, and soil nutrient availability affect the most universal indicators of quality, plant elemental composition and stoichiometry (Sterner and Elser 2002; Fan et al. 2015) and the allocation of plant secondary compounds (PSCs) that reduce nutrient availability post-ingestion (Bryant et al. 1982). Considering that biogeochemical features influence plant quality, a given browse species varies in nutrient content across individual home ranges and population ranges of herbivores (Leroux et al. 2017; Heckford et al. 2021). Despite accounting for a large portion of plant-herbivore interactions, the effects of naturally occurring intraspecific forage variation are not adequately understood.

Under the ecological stoichiometry framework, herbivores should prefer to consume plants with more available nitrogen (N) and phosphorus (P; i.e., greater N and P concentrations and lower PSC concentrations) as this increases digestive and assimilation efficiency (Elser et al. 2000; Boersma et al. 2008; González et al. 2018). Indeed, food quality impacts herbivore growth, body condition, and fitness (DeMott et al. 1998; McArt et al. 2009; Parker et al. 2009; Felton et al. 2018). Intraspecific forage choices may account for a large proportion of foraging decisions by specialists or generalists in systems of low plant diversity, but studies most often test herbivore browse selection using plant quality variation across species or age classes (but see Nie et al. 2015). Foraging studies that do investigate intraspecific forage selection often rely on fertilizer treatments rather than capturing natural variation (e.g., Ball et al. 2000), which may exaggerate any effect. If herbivore intake rate (e.g., Dostaler et al. 2011) and body condition (e.g., Rodgers and Sinclair 1997) respond to interspecific differences in forage quality, or fertilizer treatments (e.g., Schmitz et al. 1992, Ball et al. 2000), then they should also respond to intraspecific variation in forage species (Lawler et al. 1998).

While food quality is a major determinant of animal condition, studies that assess animal responses to food quality should also consider current energetic demands. Animals must acquire enough energy for body maintenance, tissue production, and reproduction through consumption (Hillebrand et al. 2009; Sperfeld et al. 2017). Energy requirements change with environmental conditions, life stages, seasons, and body conditions (Kooijman 2009). For example, endothermic animals that neither hibernate nor migrate have adapted to lower their metabolisms during winter to reduce food intake requirements and survive the resource drought (Chappel and Hudson 1978; Moen 1978). These animals use insulative winter pelage to reduce heat loss (Sheriff et al. 2009) or reduce daily energy expenditure (Humphries et al. 2005). Additionally, metabolisms rise when ambient temperatures range outside the thermal neutral zone of the endotherm (Chappel and Hudson 1978; Hillebrand et al. 2009; Sheriff et al. 2009). Animals have thus evolved flexible food intake rates in response to temporally variable energetic demands. For example, lactating mice (*Mus musculus*) have 203% higher energy demands than non-reproducing females, and throughout the lactation period, have up to 311% higher consumption rates than non-lactating females (Speakman and McQueenie 1996).

The amounts of each food nutrient or ‘currency’ an animal requires will change based on the supply and demand of said currency (Barboza et al. 2009; Simpson and Raubenheimer 2012). When a herbivore requires more of a particular currency than is available, the herbivore should be more selective for that given currency to maintain nutritional homeostasis (Villalba and Provenza 2007; Hillebrand et al. 2009). For instance, herbivores fed unnaturally high protein diets may become fibre limited and select more fibrous forage (Hodges and Sinclair 2003). When fed a diet deficient in phosphorus and calcium, lambs (*Ovis aries*) preferred supplements with those nutrients (Villalba et al. 2006). Broadening this framework: herbivores originating from habitats of low N and P availability are likely to be more nutritionally limited and thus more selective for plants N and P than conspecifics from habitats of relatively high N and P availability (Wagner et al. 2013).

Animals in the boreal forest are more nutritionally limited compared to those in more diverse, less seasonal, and more productive climates. As herbivores navigate a community of few, low-quality browse species, they should make more within-species foraging decisions. The snowshoe hare (*Lepus americanus*) is a widespread and cyclical keystone herbivore of the North American boreal and experiences highly seasonal environment (Humphries et al. 2017; Krebs et al. 2018). Hares, lower their metabolic rates to survive winter by growing dense, white coats and reducing activity levels (Sheriff et al. 2009). Hares with winter coats have lower resting metabolic rates across a wide range of ambient temperatures (−20 to 10 °C; Sheriff et al. 2009). Individual hares within a population differ in the onset of winter coat growth (Zimova et al. 2016), thus individuals with different levels of insulation can experience the same ambient temperatures, and potentially different energy expenditures. As a small herbivore, the snowshoe hare has a high and variable metabolism with low body fat storage (Sheriff et al. 2009), making it a very energetically sensitive consumer (Whittaker and Thomas 1982) and prone to weight loss in response to low food quality (Rodgers and Sinclair 1997, Ellsworth et al. 2013; see Box 1 for more details on snowshoe hare and congeneric ecology).

In Newfoundland, Canada, we sought to test if naturally occurring intraspecific variation in black spruce (*Picea mariana*) stoichiometry affected feeding choices and, consequently weight loss of snowshoe hares with different energetic demands and nutrient availabilities. Black spruce is a common item in Newfoundland snowshoe hare diets (Dodds 1960). Our study is an attempt to fill a gap in snowshoe hare foraging literature (Box 1) and test the effectiveness of plant stoichiometry as a nutritional indicator. We measured N and P compositions and PSC concentrations of black spruce (*Picea mariana*) across a 500 m by 500 m snowshoe hare trapping grid and located areas of the highest-quality (higher % N and P, lower PSC) and lowest-quality (lower % N and P, higher PSC) spruce. We then measured responses to the two qualities of spruce by snowshoe hares from the same trapping grid. To vary energetic demands, we ran cafeteria experiments in the autumn when temperatures were variable and cooling, and hares were moulting their brown summer coats and growing white winter coats. To proxy nutrient availabilities, we measured the spruce N and P compositions for the home ranges of each hare. Using these premises, we tested three non-exclusive hypotheses:

H1. The Intraspecific Choice Hypothesis: within a plant species, herbivores prefer to consume plant matter of higher quality, and those that display a stronger preference better maintain their body condition.
H2. The Energetic Demand Hypothesis: heightened energetic demands increase herbivore body maintenance costs, daily intake requirement, and demand for digestible carbon, therefore reducing the magnitude of H1 predictions.
H3. The Nutrient Availability Hypothesis: living in an environment of low-nutrient availability increases herbivore body maintenance costs and demand for limiting nutrients, therefore increasing the magnitude of H1 predictions.

We predicted that hares would prefer black spruce of higher quality, and those that more strongly exhibited this preference would lose less weight during trials (H1). Assuming that coat insulative ability and warmer temperatures decrease energy expenditure, hares with less-developed, or browner, winter coats (Sheriff et al. 2009) or those experiencing colder temperatures (Sinclair et al. 1982) would be at greater risk of weight loss, consume more total spruce, and show a lower preference for spruce N and P (H2; Simpson and Raubenheimer 2012). Lastly, hares whose ranges contained lower spruce N and P compositions would be at greater risk of weight loss and show a stronger preference for spruce N and P (H3; Hillebrand et al. 2009; Wagner et al. 2013).

## Methods

### Trapping grid and plant-snowshoe hare sampling

We carried out this work on a 25-ha snowshoe hare trapping grid located in eastern Newfoundland, Canada, during October and November of 2018 and 2019. The area typically experiences daily mean temperatures of 7.4°C (SD = 1.4) and 2.3°C (SD = 1.3) and an average monthly precipitation of 93.1 mm and 80.9 mm during October and November, respectively (Environment Canada 2019). Our trapping grid consists of planted white spruce (*Picea glauca),* which have grown above hare browsing height, and naturally occurring black spruce (*Picea mariana*) and white birch (*Betula papyrifera*). Lowland blueberry (*Vaccinium angustifolium),* Sheep laurel (*Kalmia angustifolia),* and Labrador tea (*Rhododendrum groenlandicum*) comprise the understory. In the summer, there is a herbaceous ground cover that dies off by autumn. Lowbush blueberry stems persist into autumn and are highly preferred by hares. We could not use blueberry for this study due to biomass limitations. Based on trapping records, the grid housed approximately 55 and 60 individual hares during 2018 and 2019 respectively. By tracking the locations of 27 hares on this grid using very high frequency (VHF) collars for three consecutive summers (2017-2019), we had previously found that their home range core areas (50% isopleths using kernel method) at these densities are 2.974 (± 2.351) ha on average (Rizzuto et al. 2020).

The trapping grid contains 50 Tomahawk live-traps (Tomahawk Live Trap Co. Tomahawk, WI, USA) arranged approximately 75 m apart along six transects, with traps spaced 55 and 37 m apart at the ends of transects to connect the trapping lines (Figure A1). We clipped samples of black spruce between late-June and early-August of 2016. While spruce species are not as preferred by hares compared to other species (Dodds 1960; Rodgers and Sinclair 1997), more preferred species were not as abundant in our study area. Additionally, Dodds (1960) showed that spruce species are a common diet item of snowshoe hare in Newfoundland, when highly available compared to more preferred species. Therefore, we used black spruce as our test species to sample consistently across our study area and identify intraspecific gradients of qualities. To control for effects of age class on elemental compositions, we only sampled from black spruce that were 0-2 m in height. Starting in the NW corner of an 11.3 m sample radius, we moved clockwise and collected the foliar and leaf-stem material of one individual per intercardinal direction (NW, NE, SE, and SW) until we had collected an approximate wet weight of 20 grams, and froze samples at −20°C until composition analysis. In addition, we measured two habitat variables at each trap location: the diameter at breast height (DBH; cm) of five dominant trees and the canopy closure of each intercardinal direction using a spherical crown densiometer.

We regularly trapped each spring and fall on this grid and throughout the study period to capture hares for cafeteria experiments and create a sample of known individuals. Throughout trapping, we gave hares unique ear tags upon first capture and recorded the trap location for every individual captured to determine the home range of individuals. We recorded the weight (nearest 0.02 kg), sex, right hind foot measure (mm), and age class (adult or juvenile, according to guidelines by Keith et al. 1968) for every individual captured.

### Spruce elemental and PSC analyses

We quantified black spruce elemental contents of carbon (C), nitrogen (N), and phosphorus (P) through elemental composition analysis performed at the Agriculture and Food Lab (AFL) at the University of Guelph on 10 g samples. C and N compositions were determined using an Elementar Vario Macro Cube, and P compositions were determined using a microwave acid digestion CEM MARSxpress microwave system and brought to volume using Nanopure water. The clear extract supernatant was further diluted by 10 to accurately fall within the calibration range and reduce high level analyte concentration entering the inductively coupled plasma mass spectrometry detector (ICP-MS). Black spruce samples (~4 g each) were analysed for PSC composition at the Laboratoire PhytoChemia Inc in Quebec, Canada, where the phytochemical composition was determined using a gas chromatography solvent extraction with an internal standard and a correction factor (Cachet et al. 2016). This procedure produced mg/g measures of individual terpene compounds.

### Cafeteria Offerings

To confirm that the spruce composition results were not stochastic, we performed a regression to test if spruce N, P, and PSC compositions can be explained by canopy cover and DBH, two habitat features characteristic of plant growing conditions. This test is based off the carbon-nutrient hypothesis, which predicts that plants in better growing conditions acquire more limiting nutrients, grow faster, and therefore produce less chemical defence against herbivory (Bryant et al. 1982). We predicted that higher quality spruce (high N and P, low PSC) would correlate with better growing conditions or larger DBH and canopy cover. We operated under the assumption that nearby individuals would experience the same growing condition characteristics and exhibit similar quality trends. Because spruce N, P, and PSC content did not perfectly correlate across the grid (Figure A2), we used N as our primary indicator of quality, given it is regularly considered the limiting element of terrestrial systems (White 1993). We chose six sampled trap locations to clip spruce offerings for experiments, three adjacent sites with the highest % N, and three adjacent sites with the lowest % N according to the lab analyses (Figure A1, Figure A2).

During the study, we harvested black spruce from within 15 m of the six sample locations as offerings for cafeteria experiments. We clipped twigs (< 0.3 m from the terminal end) from low branches (< 1.5 m) of adult trees (> 2 m). We clipped trees over 2 m in height to avoid exhausting juvenile trees and because adult spruce are more palatable to hares and are less likely to cause acute weight loss, based on observations using white spruce (*Picea glauca),* a congeneric of black spruce (Rodgers and Sinclair 1997). We assumed adult trees in the autumn would have similar relative elemental compositions as the juveniles (< 2 m) originally sampled in the summer (i.e., areas with higher N compositions in juveniles are also areas with higher N compositions in adults). To confirm this assumption, in 2019, we sampled approximately 17 wet grams from the adult trees we clipped at a given trap location and date and then sent these to AFL for the same elemental composition analysis as done on original samples to confirm quality ranking. Subsamples from adult trees in autumn followed similar stoichiometric trends as juvenile trees from summer, but adult trees had lower N and P compositions, which we assume was due to lignification with the growing season (Figures A3 and A4). We did not send subsamples for PSC content analysis but assume that adult spruce would exhibit spatially relative PSC trends as juvenile spruce, but in lower concentrations. We bagged spruce from each clipping location and date separately and kept them refrigerated at 0–5°C for the duration of the study. We categorised spruce from the three trap locations of highest N composition as the ‘high-quality’ offering, and spruce from the three trap locations of lowest N composition as the ‘low-quality’ offering (Table 1). To select final browse for cafeteria experiments, we eliminated any twigs or parts of twigs more than 5 mm in diameter or devoid of needles and ensured all twigs were less than or equal to 10 cm in length to fit inside feeding bins. We repeated the clipping and subsampling process when cafeteria experiments exhausted the spruce offerings.

**Table 1.**
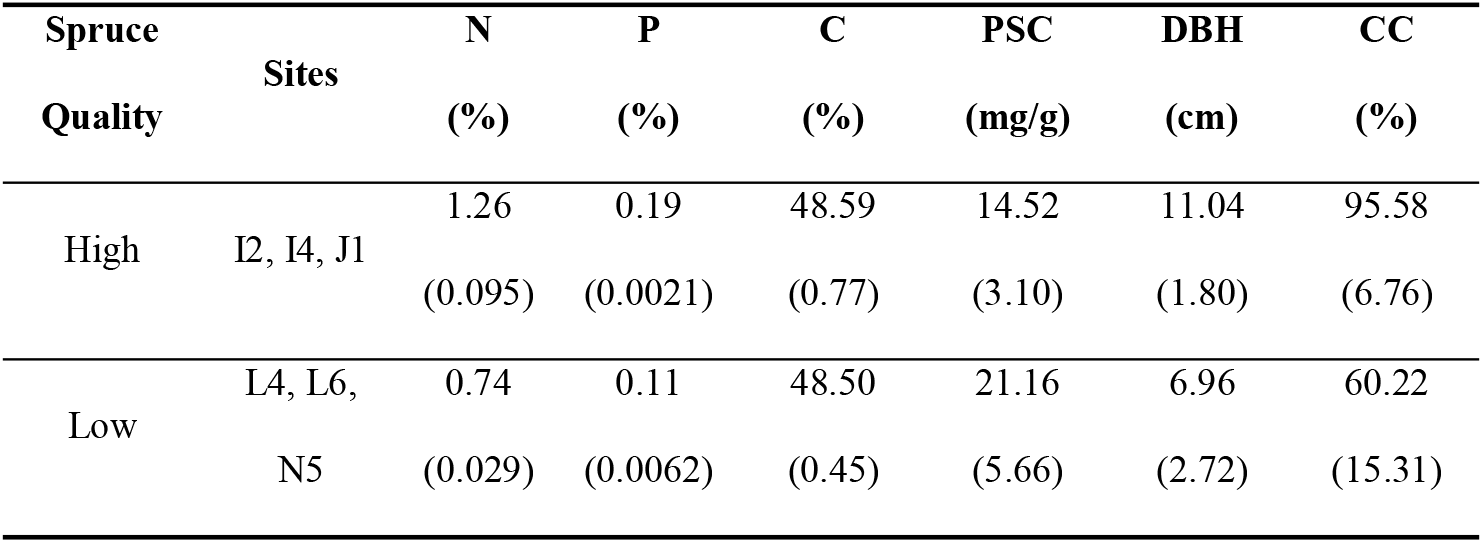
Means and standard deviations for black spruce nitrogen (N), phosphorus (P), and carbon (C) compositions (%) and plant secondary compound concentrations (PSC; mg/g), tree diameter at breast height (DBH; cm), and canopy closure (CC; %), from the trapping grid sites that provided spruce quality choices (high or low) in cafeteria experiments during 2018 and 2019. We based quality designations on nitrogen compositions measured in 2017.

### Cafeteria Experiments

All details of animal handling and experimentation were approved by Memorial University’s animal use ethics committee (AUP 18-02-EV). During October and November of 2018 and 2019, we kept individual hares for 24 h in 100 x 90 x 120 cm enclosures. Enclosures had roofs to protect hares from precipitation and a secured box for shelter (Figure A5). We placed eight enclosures in a forested area adjacent to the trapping grid and separated them by at least 10 m. We paired each enclosure with a camera trap (Reconyx Hyperfire), triggered by movement (30 s delays) to record temperatures throughout experiments (°C). We required nights in between trials to trap more hares, and therefore ran a maximum of eight cafeteria experiments per 48 h.

We captured hares overnight (< 12 hours) on two transects of the trapping grid using apple, alfalfa cubes, and timothy feed as bait (~ 100 g; hares ate all bate). We rotated from which transects we trapped hares each night to minimize repeated trapping of the same individuals. Upon checking traps, we weighed hares to the nearest 0.02 kg and only used those > 1.3 kg for experiments (Keith et al. 1968). We allowed individuals to be tested up to three times (trial numbers) per year and assumed habituation did not carry over across years. We did not use hares in consecutive cafeteria experiments (> 4 nights between trials). Experimentally-induced weight loss is a common repercussion of single species feeding trials (Rodgers and Sinclair 1997). Hares that had already undergone trials had to have recovered to within 5% of original weight to be included in other trials. We recorded trial number (1 to 3), sex, coat colour (visual estimation of % white, to the nearest 5%), and starting mass at time *t* (*Hm_t_*) The same observer assessed coat colour for all experiments.

In enclosures, we provided hares water ad libitum and approximately 130 g of high- and low-quality spruce (260 g total) in identical secured baskets on opposite sides of the enclosure (1 m apart). We randomly assigned the side of each spruce choice. Before experiments, we measured the starting mass (*SBm_t_*) of each spruce pile to the nearest 0.1 g. After 24 hours, we terminated experiments and held hares temporarily (< 1 h) to feed on timothy and apple and recover. We then reweighed hares at time *t* + 1 (*Hm*_*t*+1_) to the nearest 0.02 kg and released them at the locations of their capture. We assumed that gut content affected hares equally between initial and final weights because we always weighed hares when guts were presumably full. Finally, we weighed the spruce twigs remaining from each offering (*Sbm*_*t*+1_) to the nearest 0.1 g. We previously confirmed that overnight water loss has a negligible effect on spruce mass.

### Statistical analyses

For each experiment, we calculated the total wet mass intake, Λ, of the two piles where *i*=*h* is high-quality and *i*=1 is low quality (*I_i_* = *Sbm_t_* – *Sbm*_*t*+1_; g·day^−1^) and the percent body mass lost during trials (*Δ*Hm* = ((*Hm_t_* – *Hm*_*t*+1_)/*Hm_t_*) × 100*). We divided *I_i_* by the starting mass of the individual (*Hm_t_*) to calculate the daily intake rate (g·kg^−1^·day^−1^) for the high-quality spruce offering (*IR_h_*) and the low-quality spruce (*IR_l_*). In addition, we calculated the preference of high-quality spruce (*Pr* = *IR_h_* – *IR_l_*; g·kg^−1^ oday^−1^). To estimate nutrient availabilities for the Nutrient Availability Hypothesis, we calculated home range N and P values for each individual by averaging the black spruce N and P values across all trap locations where the individual was caught during the study period (October–November 2018, 2019). To estimate spruce elemental composition in trap locations devoid of spruce within 2 m, we interpolated results from sampled locations across the grid using ArcGIS Spatial Analyst inverse distance weighting interpolation (Childs 2004). We recorded minimum ambient temperatures from camera trap recordings of experiments. We tested for correlations (Pearson’s r) between coat colour and temperature, and home range N and P, to confirm there was no interaction between nutrient availability and energetic demand. We observed that hares ate significantly more spruce when habituated to experiments (second or third trial) than when non-habituated (*t* = 2.16, *p* = 0.034; Figure A6). The increase in intake rate with habituation did not translate to a significant difference in spruce preference (t = 0.60, p = 0.55; Figure A6) or weight loss (t = −0.287, p = 0.775; Figure A6). Therefore, we accounted for habituation when testing our predictions for feeding responses.

Our three, non-exclusive hypotheses have two components–feeding responses and body condition responses to spruce quality choices, energetic demands, and nutrient availabilities. We tested the predictions with two sets of model comparisons, one per response, using second-order Akaike Information Criterion (AICc) in program R (Version 3.6.3; R Development Core Team 2018), using the ‘aictab’ function from the *AICcmodavg* package (Mazerolle 2017). We begin by outlining models used to test the feeding component of our predictions. We compared nine linear mixed models (Table A1) that all measured the effect of spruce quality on the intake rate of a pile (*IR_h_* or *IR_I_*) to test the Intraspecific Choice Hypothesis on data paired by experiment (random effect = ‘experiment ID’). All models included habituation (binary) as a fixed-effect (see Table A3 for results and discussion when using only non-habituated data). The models compared included: only spruce quality as a direct test of the Intraspecific Choice Hypothesis (‘Base’), coat colour and minimum temperature as a test of the Energetic Demand Hypothesis (‘Energetic’), home range N and P as a test of the Nutrient Availability Hypothesis (‘Nutrient’). Additionally, we tested each variable independently (‘Coat Colour’, ‘Temperature’, ‘Nitrogen’, ‘Phosphorus’), and created a ‘Full’ model with all competing variables and an intercept only, ‘Null’ model (Table A1). All models included the interaction between each energetic or nutritional variable and spruce quality (Table A1). Beta-coefficients from non-interacting effects show effects on overall consumption rates and Beta-coefficients from interacting effects show effects on preference between spruce qualities (positive = a positive effect on preference for high-quality spruce).

Next, we tested the body condition component of our predictions, which paralleled those of the feeding component. We compared nine linear models that all measured the effect of preference for high quality spruce (*P*) and total intake rate (*IR*) on mass lost during trials (*m_i_*) to test the Intraspecific Choice Hypothesis. We compared tests for all three hypotheses similar to as done for feeding responses. The models include: ‘Base’, ‘Energetic’, ‘Nutrient’, ‘Coat Colour’, ‘Temperature’, ‘Nitrogen’, ‘Phosphorus’, and the ‘Null’ (Table A1). All models included the interaction between each energetic or nutritional variable and preference (Table A1).

## Results

### Black spruce and habitat measures

Of the 50 sampling sites on our trapping grid, 36 had black spruce present and 14 required interpolated spruce elemental composition values. From sampled spruce (n = 36), the mean black spruce C, N, P, and PSC compositions were 49.03 ± 0.70%, 1.01 ± 0.18%, 0.14 ± 0.036%, and 17.20 ± 4.95 g/mg, respectively. The mean canopy closure and DBH for all 50 trap locations were 77.84 ± 28.05% and 9.16 ± 2.28 cm respectively. N significantly correlated positively with P compositions (t = 5.08, p < 0.001; Figure A2) and negatively with PSC (terpenes) compositions (t = −2.60, p = 0.014; Figure A2). Phosphorus negatively correlated with PSCs, but not significantly (t = −1.46, p = 0.15; Figure A2). Spruce elemental and phytochemical compositions were not stochastic because canopy closure and DBH combined explained 35.5%, 25.1%, and 15.2% of spruce N, P and PSC variation across the grid, respectively (multiple R^2^; Figure A7). Canopy closure had a more significant effect towards N (t = 3.094, p < 0.01), P (t = 3.041, p = 0.0046), and PSC (t = −1.64, p = 0.11) than DBH (N t = 0.931, p = 0.35; P t = −0.402, p = 0.69; PSC t = −1.64, p-value = 0.11). Canopy closure and DBH did not explain variation in spruce C compositions (R^2^ = 0.01). We clipped high-quality spruce for cafeteria experiments where N compositions ranged from 1.17 – 1.39%, P ranged from 0.169 – 0.197%, and PSC ranged from 11.36 – 17.57 mg/g (Table 1). We clipped low-quality spruce offering where N compositions ranged from 0.71 % – 0.76 %, P ranged from 0.101 % – 0.113 %, and PSC ranged from 14.89 – 25.88 mg/g (Table 1). We confirmed that subsampled clippings from high-quality sites had more N and P than low-quality sites (Figures A2 and A3).

### Cafeteria Experiments

We conducted a total of 75 cafeteria experiments in October and November of 2018 (n = 22) and 2019 (n = 53). Our sample included 48 individuals (M = 20; F = 28), five of which were tested in both years. In two experiments, hares escaped before we could reweigh them. There were 51 experiments with non-habituated hares (first trials) and 24 with habituated hares (second and third trials). Individuals began experiments weighing an average of 1500 g and lost a mean of 5.8 ± 2.73 % body mass during experiments. Minimum temperatures during experiments ranged from −4°C to 6°C (median = 0). Hares ranged from 0 to 90% white, and the median coat colour was 10% white. The mean individual home range spruce N was 1.01 ± 0.16% and, mean range P was 0.14 ± 0.031%. Energetic variables were not correlated with nutritional variables (−0.05 < r < 0.14) and, therefore, did not affect out hypothesis tests. Minimum ambient temperatures weakly correlated (r = −0.55) with whiter coats, as expected with this seasonal transition.

Placement of spruce choices, relative to enclosure entrance and shelter, did not influence intake rates (t = 1.463, p = 0.15). Hares sorted through spruce piles, spreading twigs around within a 0.25 m radius of the baskets. Hares tended to reject the last 3-5 mm of spruce terminal ends, as these tips of twigs frequently remained post experiment. Hares ate on average 125.35 ± 35.9 g/day (*I_i_*) or 82.98 ± 22.8 g·kg^-1^·day^−1^ (*IR_i_*) of spruce during experiments, 43.65 ± 15.5 g·kg^-1^·day^−1^ from high quality spruce (*IR_h_*) and 39.32 ± 14.9 g·kg^−1^·day^−1^ from low quality spruce (*IR_l_*).

### eeding response

Overall, spruce quality alone could not predict snowshoe hare feeding responses during cafeteria experiments (Base model; Table A2). Energetic and Temperature models parsimoniously explained intake rates and quality preference (Table A2). All other models had ΔAICc greater than 2 (Table A2). Based on the Energetic model, minimum ambient temperature (°C; β = −1.73 ± 0.61, t = −2.83, p = 0.007) and coat colour (% white; β = −12.70 ± 6.82, t = −1.86, p = 0.07) negatively correlated with spruce intake rate (Table 2; Figure 2), while spruce quality and habituation had no significant effect. According to the interaction coefficients, hares that had whiter coats ate more high-quality than low-quality spruce compared to hares with browner coats (β = 19.61 ± 9.25, t = 2.12, p = 0.042). Similarly, though not significantly, hares in warmer temperatures preferred high quality spruce more than hares in colder temperatures (β = 0.96 ± 0.83, t = 1.16, p = 0.26; Table 2; Figure 2). In the second-top ranked Temperature model, we observed a similar trend between minimum temperature and total intake rate (β = −1.11 ± 0.52, t = −2.12, p = 0.041). In the Temperature model, the interaction between temperature and spruce quality was near-zero (p = 0.99; Table 2).

**Figure 1.**
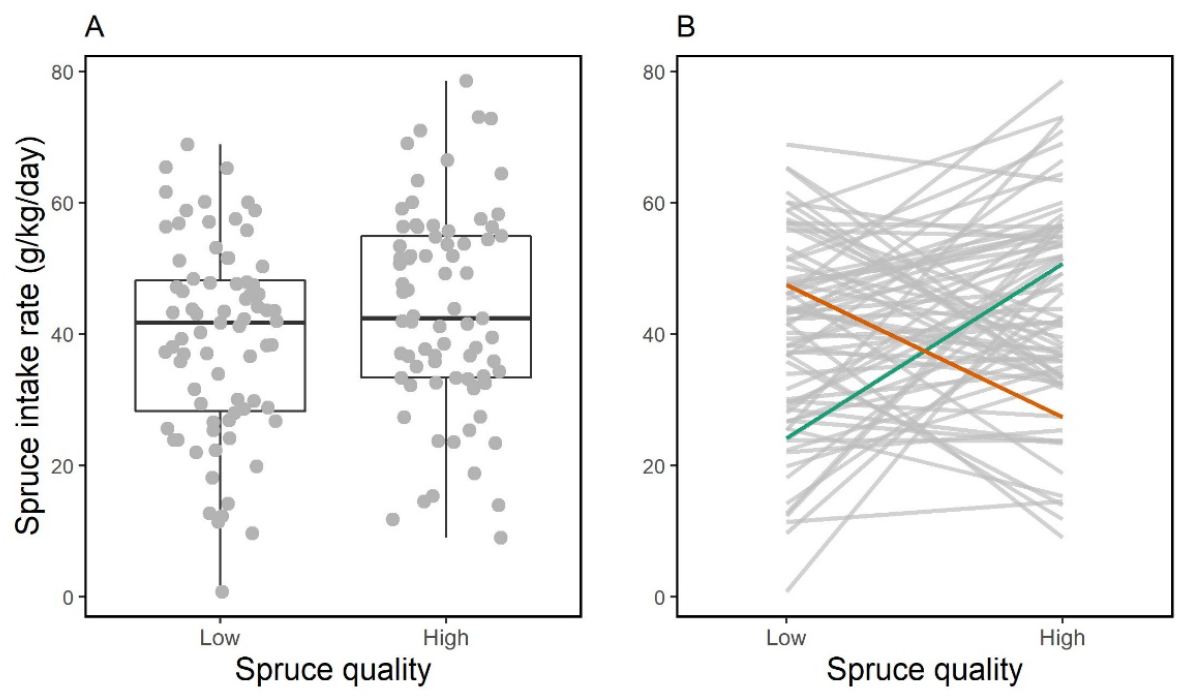
A) Overall trend and B) individual trends of snowshoe hare consumption (g·kg^−1^·day^−1^) of browse piles when offered two nutritional ranks of black spruce during 24-hour cafeteria experiments (75 experiments, 150 observations). In panel B results from two experiments that yielded opposing feeding preferences are highlighted. The green line shows an individual that preferred the high-quality spruce (positive *P*), and the red line shows an individual that preferred the low-quality spruce (negative *P*).

**Figure 2.**
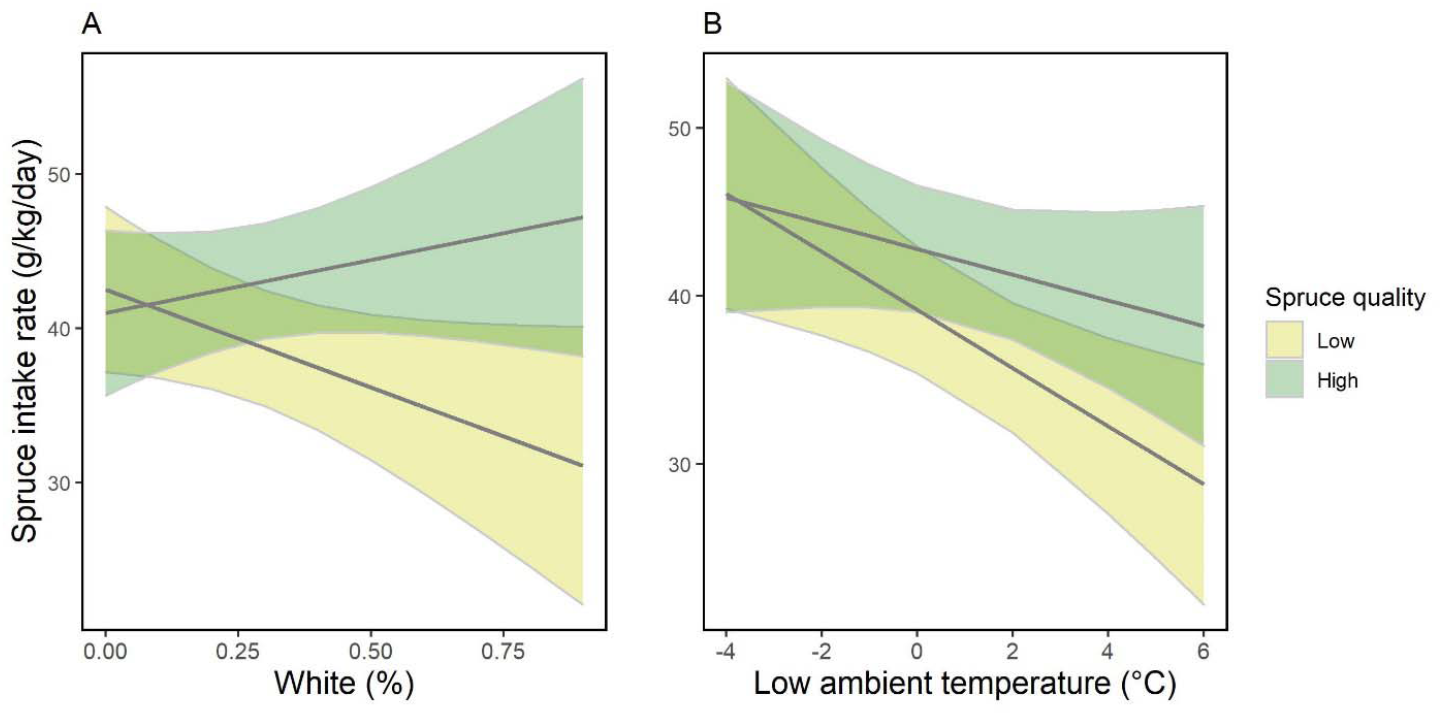
Linear relationships with 0.95 confidence intervals (bands) between intake rate (IR; g·kg^−1^·day^−1^) of each spruce offering (low quality = yellow, high quality = green) by individual, snowshoe hares during cafeteria experiments (75 experiments, 150 observations) in relation to A) individual coat insulation (% white) and B) minimum ambient temperature during experimentation (Energetic-intake model).

**Table 2.** Fixed-effect coefficients (standard errors) and marginal and conditional R^2^s from all AICc compared linear mixed models (paired by experiment) predicting snowshoe hare intake rate (g·kg^−1^·day^−1^) of black spruce high and low qualities when offered in pairs during individual cafeteria experiments (150 observations, 75 experiments). Top-ranked models by AICc are bolded. Note: *p<0.1, **p<0.05, ***p<0.01.

| Model | Model Coefficients | | | | | | | | | | $R^2_m$ | $R^2_c$ |
| --- | --- | --- | --- | --- | --- | --- | --- | --- | --- | --- | --- | --- |
|  | Hab. | Qual. | Temp | Temp*<br>Qual | Coat | Coat* Qual | N | N* Qual | P | P* Qual |  |  |
| Null | 5.96**<br>$\pm 2.72$ | | | | | | | | | | 0.03 | 0.11 |
| Base | 5.96**<br>$\pm 2.72$ | 4.32* ±<br>2.3 | | | | | | | | | 0.05 | 0.15 |
| **Temp** | **4.17 ±**<br>**2.65** | **4.32\* ±**<br>**2.36** | **-1.11\*\* ±**<br>**0.52** | **0 ±**<br>**0.71** |  |  |  |  |  |  | **0.11** | **0.15** |
| Coat | 5.34*<br>$\pm 2.77$ | 0.73 ±<br>3.05 | | | -2.42 ±<br>5.93 | 13.67* ±<br>7.79 | | | | | 0.08 | 0.18 |
| **Energetic** | **4.36 ±**<br>**2.66** | **-1.52 ±**<br>**3.59** | **-1.73\*\*\***<br>**$\pm 0.61$** | **0.97 ±**<br>**0.83** | **-12.7\***<br>**$\pm 6.82$** | **19.61\*\* ±**<br>**9.25** |  |  |  |  | **0.13** | **0.19** |
| Nitrogen | 6.42**<br>$\pm 2.71$ | 5.3 ±<br>20.42 | | | | | -14.67<br>$\pm 14.81$ | -0.96 ±<br>19.95 | | | 0.07 | 0.15 |
| Phosphorus | 6.07**<br>$\pm 2.72$ | -8.64 ±<br>13.3 | | | | | | | -75.9 ±<br>69.08 | 91.51 ±<br>92.47 | 0.06 | 0.16 |
| Nutrient | 6.45**<br>$\pm 2.7$ | 7.04 ±<br>20.18 | | | | | -6.4 ±<br>20.38 | -28.06 ±<br>27.32 | -55.69 ±<br>94.83 | 182.31 ±<br>127.46 | 0.08 | 0.17 |
| Full | 4.73*<br>$\pm 2.68$ | 7.08 ±<br>19.77 | -1.67***<br>$\pm 0.62$ | 1.03 ±<br>0.84 | -12.4*<br>$\pm 6.8$ | 19.96** ±<br>9.21 | 3.04 ±<br>19.87 | -33.71 ±<br>26.83 | -56.65 ±<br>91.16 | 180.38 ±<br>123.65 | 0.15 | 0.22 |

\*Temp = Temperature; Coat = Coat Colour; Hab. = Hare habituation; Qual = Spruce quality; N = Nitrogen; P = Phosphorus

### Weight loss responses

Conversely, the top-ranked model explaining hare weight loss was the Base model (Table A2). All other models had ΔAICc greater than 2.00 (Table A2). We observed, in the Base model, total intake rate to not significantly affect weight loss during experiments (β = −0.021 ± 0.014, t = −1.47, p = 0.14; Table 3). However, the preference for high-quality spruce was significantly (t = −2.43; p = 0.018) correlated to lower weight loss (β = −0.037 ± 0.015; Table 3; Figure 3).

**Table 3.**
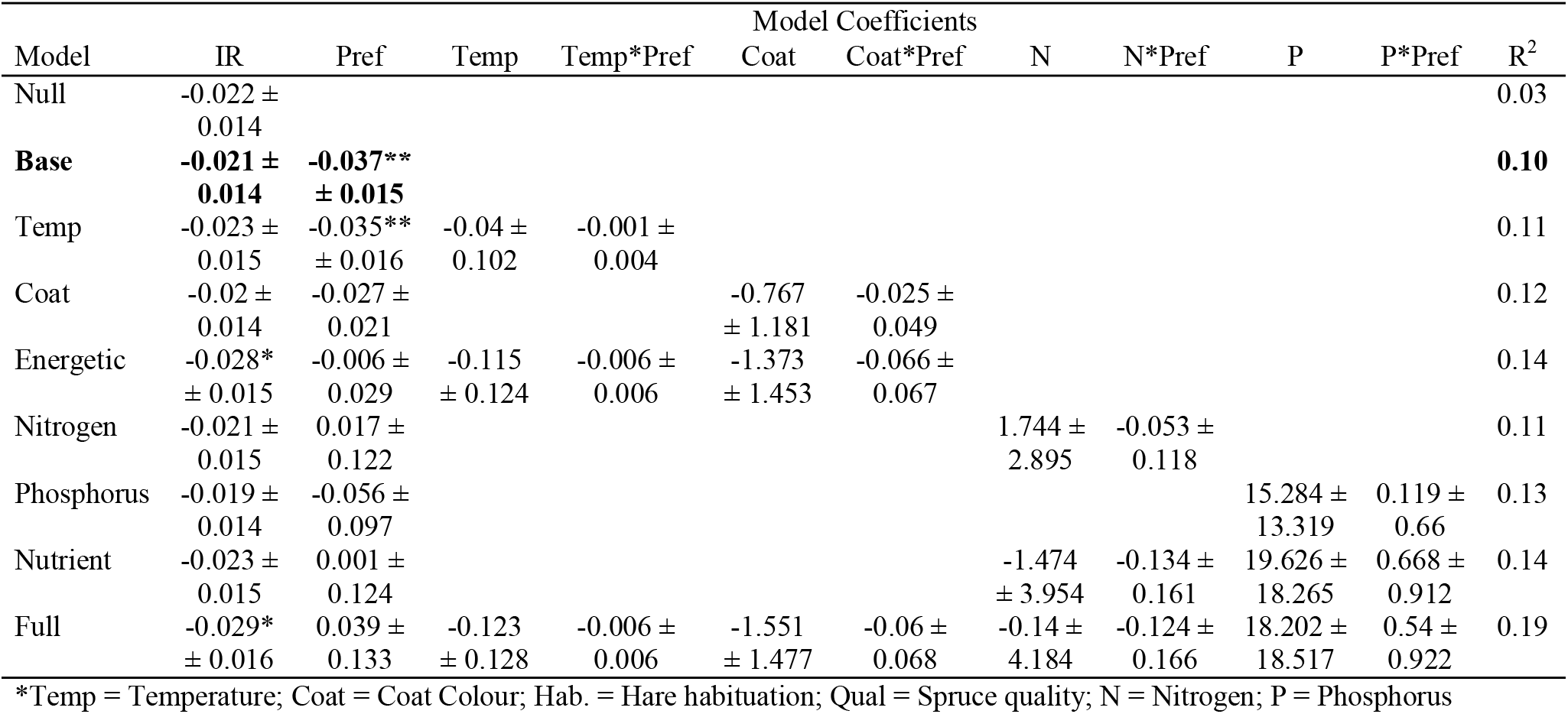
Fixed-effect coefficients (standard errors) and R^2^s from all linear models compared using AICc to predict snowshoe hare weight loss (% per day) during cafeteria experiments that tested intraspecific preferences of spruce (n = 73). Top-ranked models by AICc are bolded. Note: *p<0.1, **p<0.05, ***p<0.01.

**Figure 3.**
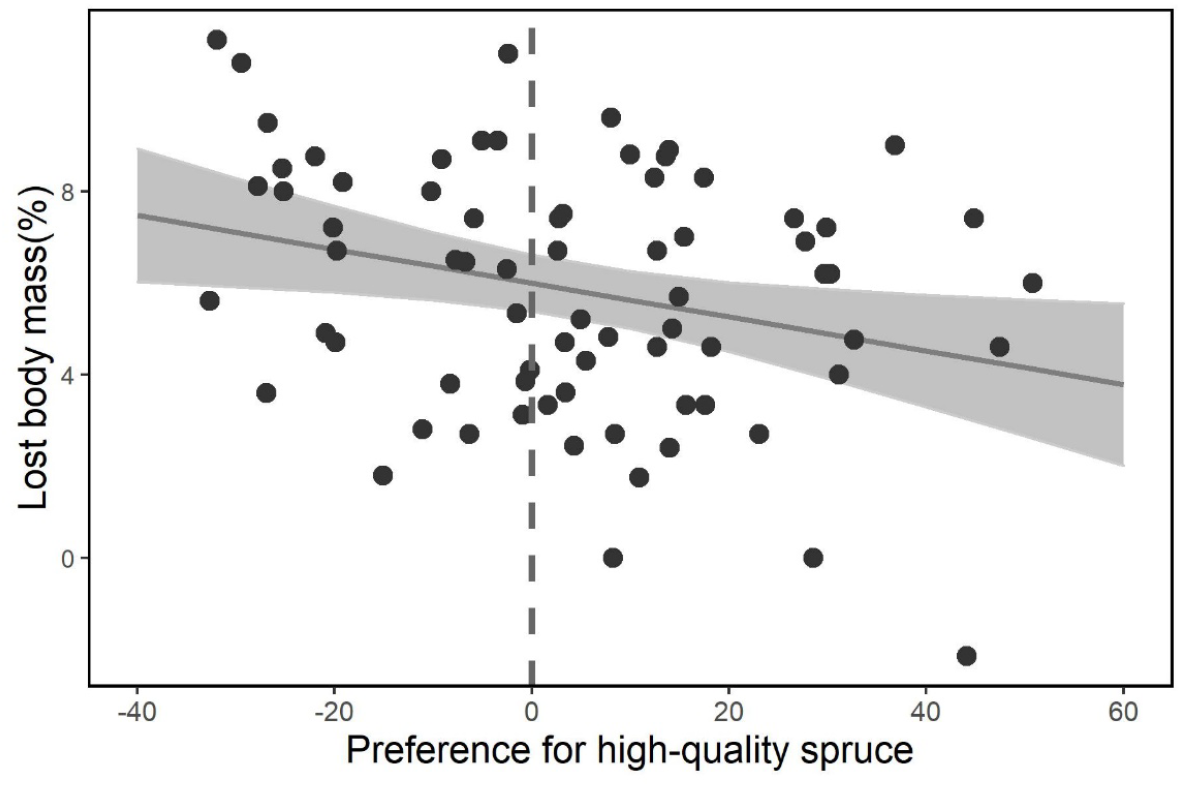
Body mass change (% lost per day) by snowshoe hares during cafeteria experiments, which tested their feeding response to two qualities of black spruce, in relation to their preference for high-quality spruce (high-quality intake rate – low quality intake rate; g·kg^−1^day^−1^). The regression line was taken from the Base-weight loss model (R^2^ = 0.11).

## Discussion

Herbivore feeding choices and body conditions vary in response to available plant qualities and limit body energetic and nutritional demands (Barboza et al. 2009). We conducted cafeteria-style experiments using a boreal herbivore, the snowshoe hare, and its locally available forage. Capturing the natural variation in intraspecific forage and capitalizing on changing energetic demands, we tested non-exclusive hypotheses on forage choice and its implications for weight loss. Specifically, we tested if hares differentiate between two qualities of black spruce, how this translated to weight loss during experiments, and if these responses are affected by energetic and nutritional demands. Overall preference for higher quality spruce was driven by energetic demands, but irrespective of energetic state, greater selection for higher quality spruce led to less weight loss. We did not find any effect by our proxies for nutrient availability on either feeding or body condition responses. Our findings suggest that plant growing conditions ultimately affect herbivore body conditions, and that herbivore feeding choices depend on energetic state (Figure A8).

The Intraspecific Choice Hypothesis (H1) assumes that herbivores should browse the most digestible and high-quality food, and not doing so could hinder body condition and fitness (Parker et al. 2009; Wam et al. 2018). Overall, preference for high-quality spruce, when not considering any other factors, was insignificant (Figure 1). Weight loss was influenced by feeding preference (Figure 3), as predicted. However, weight loss was not influenced by other variables such as total intake rate or temperature. We illustrate that the individuals with the strongest preference for high-quality spruce (+40 g·kg^−1^·day^−1^) lost on average 2.2% less weight than hares with the strongest, opposing preference for low-quality spruce (−20 g·kg^−1^·day^−1^). Other feeding experiments using hares have linked weight loss to browse quality. Quality can be indicated by many currencies including energy, protein, or PSCs, all of which influence the ability of herbivores to meet maintenance requirements (Pehrson 1984; Rodgers and Sinclair 1997; Ellsworth et al. 2013). While no single dietary currency can predict herbivore body condition (Parker et al. 2009; Raubenheimer et al. 2009; Felton et al. 2018), protein, an N-dense compound, is often a limiting currency for herbivores (Sinclair et al. 1982). Hares that preferred higher-quality spruce likely assimilated more protein, better met maintenance requirements, and hence lost less weight. Although hares did not show overall strong intraspecific preference, spruce quality influenced feeding responses when it interacted with energetic variables.

The Energetic Demands Hypothesis (H2) posits first that higher energetic needs translate into higher intake rates by the herbivore: our findings support this idea. Hares had higher intake rates at lower ambient temperatures (p < 0.01) and when sporting less insulative coats (p < 0.1). Specifically, our Energetic model indicated that hares with fully white coats eat approximately 25.3 g·kg^−1^·day^−1^ less spruce than hares with fully brown coats and that hares experiencing a minimum ambient temperature of 4°C will consume 27.68 g·kg^−1^·day^−1^ less spruce compared to those experiencing that of −4°C (Table 2). Higher intake rates in colder conditions are expected given temperature is associated with feeding rates (Sinclair et al. 1982; Camp et al. 2018). We assume the mechanism behind the coat colour-feeding relationship is metabolic change. Temperate mammals lower their metabolisms and food intake in winter with the aid of more insulative pelage (Chappel and Hudson 1978), but our work extends this to identify feeding rates across a gradient of a seasonal pelage transition. Hares with fully developed winter coats have lower resting metabolic rates than those with autumn coats (Sheriff et al. 2009). While Sheriff et al. (2009) found a binary metabolic difference, according to our results, the relationship between coat colour and metabolism may be continuous during the seasonal transition if we assume feeding rate is proportional to metabolic rate.

The Energetic Demand Hypothesis also predicts that energetically driven higher intake rates lower herbivore selective ability. Our findings supported this prediction because whiter hares had significantly stronger preferences for high-quality spruce (Figure 2). Snowshoe hares in warmer temperatures followed the same trend, though not significantly (Figure 2). While we had predicted that total intake rate would limit hare feeding preferences and drive this response, intake rate alone did not influence preference (Figure A9). However, energy budgets may influence feeding preferences independent of feeding rate. For example, Camp et al. (2018) found that in lower ambient temperatures, rabbits incorporated more fibre in their diet and assumed this was because higher fibre diets produced more heat during digestion than lower fibre diets (Chappell et al. 1997). Regardless of the mechanism, herbivores interact with plant communities differently depending on energetic states. We found hares to choose different diets depending on daily temperature variation and their stage of seasonal transition.

The Nutrient Availability Hypothesis (H3) predicts that higher nutritional requirements drive herbivores to greater preference for higher-quality browse. We did not find support for this prediction using our proxies of nutrient availabilities, i.e., spruce N and P of individual home ranges. Nutrient models were not top ranked and we did not find a significant effect of range N or P on feeding rates, food preferences, or weight loss. Our spruce quality measures were indicative of growing conditions given DBH and canopy closure, habitat measures that accounted for other tree species, explained 36.0%, 25.1%, and 26.5% of spruce N, P, and PSC variation respectively (Figure A7). While we did not detect an effect of home range nutrient availability on individual responses during experiments, the feeding decisions between spruce qualities from this same area greatly influenced hare body conditions. Longer-term, fitness effects of plant quality may only occur at larger scales where micro-climates, ecotones, rock-beds, soil types, and plant communities are more likely to vary and create wider variation in nutrient availability (McArt et al. 2009).

Plants are limited by nutrient availability and growing conditions, and herbivores are limited by plant abundance and quality. However, the effects of natural intraspecific plant quality variation on herbivore feeding and body condition remain underexplored, especially with consideration for other factors that affect herbivores, e.g., energetic and nutritional demands. Borrowing from principles of ecological stoichiometry and measuring the elemental composition of food, we addressed the multi-faceted nature of plant-herbivore interactions. We linked growing conditions to intraspecific variation in plant quality (e.g., % N) across our snowshoe hare trapping grid and found this variation to significantly affect hare body conditions during feeding experiments. Overall, feeding preferences for high-quality spruce were weak but exacerbated by individual energetic states. Feeding preferences, not energetic states, were linked to weight loss during experiments, indicating a complex interaction between energetics and feeding on body condition. In this case, energetic states influenced body condition, but only indirectly via feeding choices. Collectively our findings address the complex feedbacks between plant growing conditions, herbivory, and animal body condition. Biogeochemistry, seasons, and temperature influence plant productivity and quality (Pastor and Post 1986; Tang et al. 2018) and thus the food quality for herbivores.

## Acknowledgements

Thank you to Ally Menzies, Yasmine Majchrzak, and Michael Peers for giving insight on the field methods for this study. We thank Kara Gerrow, Julie Turner, Katrien Kingdon, Jaclyn Aubin, Alec Robitaille for help with data collection.

## Data and code accessibility statement

All data and code associated with this research are available on github.com: https://github.com/jballuffi/StoichiometryCafeteriaExperiments

## Ethics Approval

All applicable institutional and/or national guidelines for the care and use of animals were followed. The Animal Care Committee of Memorial University of Newfoundland approved our livetrapping, handling, and experimentation protocol with permit #18-02-EV.

## Consent to Participate

Not applicable.

## Consent for Publication

Not applicable.

## Oecologia Highlighted Student Paper

We would like for this paper to be considered for the Oecologia Highlighted Student Paper honor because the study’s design and findings represent novel contributions to the field of foraging ecology. Firstly, this paper is a unique example of how to conduct feeding experiments on a wild mammal population, eliminating the need for maintaining a captive sample of animals, a resource intensive practice. Secondly, this study reports significant effects of natural intraspecific variation in plant stoichiometry, an often-overlooked relationship, showing that not all plants within a species are the same to herbivores. Thirdly, the study considers energetic states when testing for foraging responses and finds that plant-herbivore interactions are dynamic relationships, influenced by the abiotic environment.

## Authors’ Contributions

JBF, EVW, SJL, YFW, MR, TRH, and ICR devised the study, JBF, EVW, SJL, YFW, MR, TRH, ICR, and JLK collected and analysed the data, JBF, EVW, SJL, YFW, MR, TRH, ICR, and JLK interpreted results, JBF led writing of the manuscript, JBF, EVW, SJL, YFW, MR, TRH, ICR, and JLK revised and edited the manuscript

