## Appendices for "Integrating plant stoichiometry and feeding experiments: state-dependent forage choice and its implications on body mass"

Total objects: 13

Box 1. Summary of snowshoe hare and congenerics feeding ecology literature.

Over recent decades the snowshoe hare has been subject to dietary studies attempting to investigate the chemical basis of their food choices (Sinclair and Smith 1984; Ellsworth et al. 2013) and frequent investigations have shown the general preference ranking of browse species for hares (Dodds 1960; Bryant and Kuropat 1980; Rodgers and Sinclair 1997). These preferences are often attributed to hare selection for protein and energy content (Rodgers and Sinclair 1997; Ellsworth et al. 2013), and avoidance of plant secondary compounds (PSCs; Bryant et al. 1985). However, findings on snowshoe hare diet selection can be more complex. In multiple cases, feeding strategies appear to balance intake rates of fibre and protein (Hodges and Sinclair 2003) or protein and secondary compounds (Schmitz et al. 1992), and findings on avoidance of PSCs confound one another (Sinclair and Smith 1984; Bryant et al. 1985). Sinclair and Smith (1984) originally speculated that the inconsistent results across snowshoe hare feeding studies were due to specific differences in defence chemicals across “species, growth stages, or even individuals”. Investigating the preference of snowshoe hare feeding within a single browse species could potentially clarify drivers of interspecific choice. Hares often prefer to browse on older trees (Bryant et al. 1985), and reject certain plant parts such as the green foliar buds of white birch (*Betula papyrifera*), green alder (*Alnus viridis*), and balsam poplar (*Populus balsamifera*; Bryant and Kuropat 1980). Laitinen et al. (2002), which tested mountain hare preference for clones of *Betula pendula* grown in different habitats*,* found tree genotype to explain much variation of feeding. However, generally, preference for intraspecific nutritional variation due to natural growing conditions remains underexplored.

Table A1. Design of linear mixed models compared to explain the intake rate (IR_i_; g·kg^-1^·day^-1^) of black spruce and linear models compared to explain weight loss (ΔHm; % lost per day) of snowshoe hares in cafeteria experiments. Both sets of model comparisons test the same combinations of energetic (temperature, temp, and coat colour, coat) and nutritional (range N and P) effects. Bolded models incorporate all independent variables relative to a hypothesis we test. All feeding response models compare the intake rate of spruce offerings within an experiment by using experiment ID as a random effect and account for habituation (Hab) by using it as a non-interacting fixed effect (e.g., IR_i_ ~ Hab + Quality is the base model for feeding response comparisons). All body condition models test the effect of preference (Pref) for high-quality spruce on weight loss (ΔHm) during experiments and account for total spruce intake rate (IR) by using it as a non-interacting fixed effect (e.g., ΔHm ~ IR + Pref is the base model for body condition response comparisons).

| *Model name* | *Feeding response models*  *Independent variable = IR_i_ (*g·kg^-1^·day^-1^*)* | *Body condition response models*  *Independent variable = ΔHm (%/day)* |
| --- | --- | --- |
| Null | Hab | IR |
| **Base** | **Hab + Quality** | **IR + Pref** |
| Coat | Hab + Quality*Coat | IR + Pref*Coat |
| Temp | Hab + Quality*Temp | IR + Pref*Temp |
| **Energetic** | **Hab + Quality*Temp + Q*Coat)** | **IR + Pref*Temp + Pref*Coat** |
| N | Hab + Quality*N | IR + Pref*N |
| P | Hab + Quality*P | IR + Pref*P |
| **Nutrient** | **Hab + Quality*N + Quality*P** | **IR + Pref*N + Pref*P** |
| Full | Hab + Quality*Temp + Quality*Coat + Quality*N + Quality*P | IR + Pref*Temp + Pref*Coat + Pref*N + Pref*P |

Table A2. AICc comparisons of all models predicting snowshoe hare feeding responses (left) and body condition responses (right) when offered two qualities of black spruce in individual cafeteria experiments. The feeding response AICc compares linear mixed models (paired by experiment) that predict snowshoe hare intake rate of the high and low-quality black spruce (n = 150). The body condition response AICc compares linear models that predict snowshoe hare weight loss during the same experiments (n = 73). Models ranked within 2 ΔAICc of the top model are bolded. For the design of all models compared, see Table A1.

| **Feeding response models** | | | | | | **Body condition response models** | | | | | |
| --- | --- | --- | --- | --- | --- | --- | --- | --- | --- | --- | --- |
| Model | K | AICc | ΔAICc | AICcWt | LL | Model | K | AICc | ΔAICc | AICcWt | LL |
| **Energetic** | **9** | **1240.54** | **0** | **0.42** | **-610.624** | **Base** | **4** | **-319.35** | **0** | **0.53** | **163.97** |
| **Temp** | **7** | **1240.73** | **0.20** | **0.38** | **-612.972** | P | 6 | -316.50 | 2.84 | 0.13 | 164.89 |
| Base | 5 | 1244.34 | 3.80 | 0.062 | -616.961 | Coat | 6 | -315.75 | 3.60 | 0.088 | 164.51 |
| Coat | 7 | 1244.73 | 4.19 | 0.051 | -614.968 | Null | 3 | -315.69 | 3.65 | 0.086 | 161.021 |
| Null | 4 | 1245.65 | 5.11 | 0.033 | -618.685 | N | 6 | -315.12 | 4.23 | 0.064 | 164.20 |
| Full | 13 | 1246.79 | 6.25 | 0.018 | -609.054 | Temp | 6 | -314.99 | 4.36 | 0.060 | 164.13 |
| N | 7 | 1246.82 | 6.28 | 0.018 | -616.014 | Energetic | 8 | -313.17 | 6.18 | 0.024 | 165.71 |
| P | 7 | 1247.39 | 6.86 | 0.014 | -616.302 | Nutrient | 8 | -312.70 | 6.65 | 0.019 | 165.47 |
| Nutrient | 9 | 1249.04 | 8.51 | 0.006 | -614.878 | Full | 12 | -305.40 | 13.94 | 0.00050 | 167.30 |

Table A3. Results from an AICc comparison that tested whether results from Table A2 would be similar when sub-setting the cafeteria experiments to only include first trials (or non-habituated hares) and therefore removing habituation as a fixed effect, like in the model designs shown in Table A1. The feeding response AICc compares linear mixed models (paired by experiment) that predict snowshoe hare intake rate of the high and low-quality black spruce (n = 102). The body condition response AICc compares linear models that predict snowshoe hare weight loss during the same experiments (n = 51). Models ranked within 2 ΔAICc of the top model are bolded. We decided to account for habituation in our models because we found hares ate significantly more spruce during their second and third cafeteria experiments (Figure A6). Results here suggest that our decision to use habituation as a fixed effect in the main analysis did not skew results because the top-ranked models on non-habituated experiments here are similar to our main results (Table A2).

| **Feeding response models** | | | | | | **Body condition response models** | | | | | |
| --- | --- | --- | --- | --- | --- | --- | --- | --- | --- | --- | --- |
| Model | K | AICc | ΔAICc | AICcWt | LL | Model | K | AICc | ΔAICc | AICcWt | LL |
| **Energetic** | **8** | **834.049** | **0** | **0.903** | **-408.25** | **Null** | **3** | **246.052** | **0** | **0.429** | **-119.771** |
| Temp | 6 | 839.849 | 5.799 | 0.05 | -413.482 | **Base** | **4** | **246.427** | **0.375** | **0.356** | **-118.779** |
| Null | 3 | 842.115 | 8.066 | 0.016 | -417.935 | Temp | 6 | 249.482 | 3.43 | 0.077 | -117.786 |
| Base | 4 | 842.907 | 8.857 | 0.011 | -417.247 | Nitrogen | 6 | 250.299 | 4.247 | 0.051 | -118.195 |
| Full | 12 | 843.034 | 8.985 | 0.01 | -407.764 | Phosphorus | 6 | 251.056 | 5.004 | 0.035 | -118.574 |
| Coat | 6 | 844.504 | 10.455 | 0.005 | -415.81 | Coat | 6 | 251.43 | 5.378 | 0.029 | -118.76 |
| Nitrogen | 6 | 845.868 | 11.819 | 0.002 | -416.492 | Energetic | 8 | 252.625 | 6.573 | 0.016 | -116.598 |
| Phosphorus | 6 | 846.06 | 12.01 | 0.002 | -416.588 | Nutrient | 8 | 254.453 | 8.401 | 0.006 | -117.512 |
| Nutrient | 8 | 849.61 | 15.56 | 0 | -416.031 | Full | 12 | 262.601 | 16.549 | 0 | -115.195 |

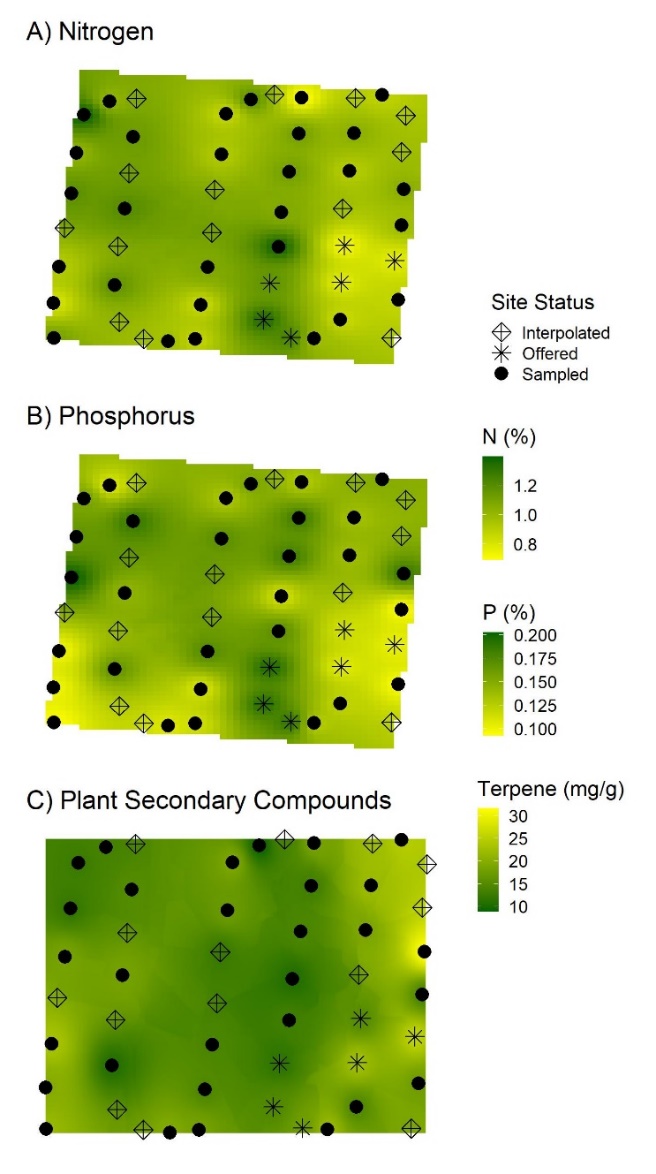

Figure A1. Interpolated maps of the trapping grid’s black spruce A) nitrogen compositions (%), B) phosphorus compositions (%), and C) plant terpene concentrations (PSCs; mg/g). Each sample location is a snowshoe hare trap location. Spruce was sampled during the summer of 2017 for N and P analysis at sites where the species was present (solid circles and stars), and these results were then used to interpolate the whole grid and give values to locations which did not have spruce within their sample plots (crosses). Areas of highest and lowest nitrogen compositions were offered (stars) in cafeteria experiments during the fall of 2018 and 2019.

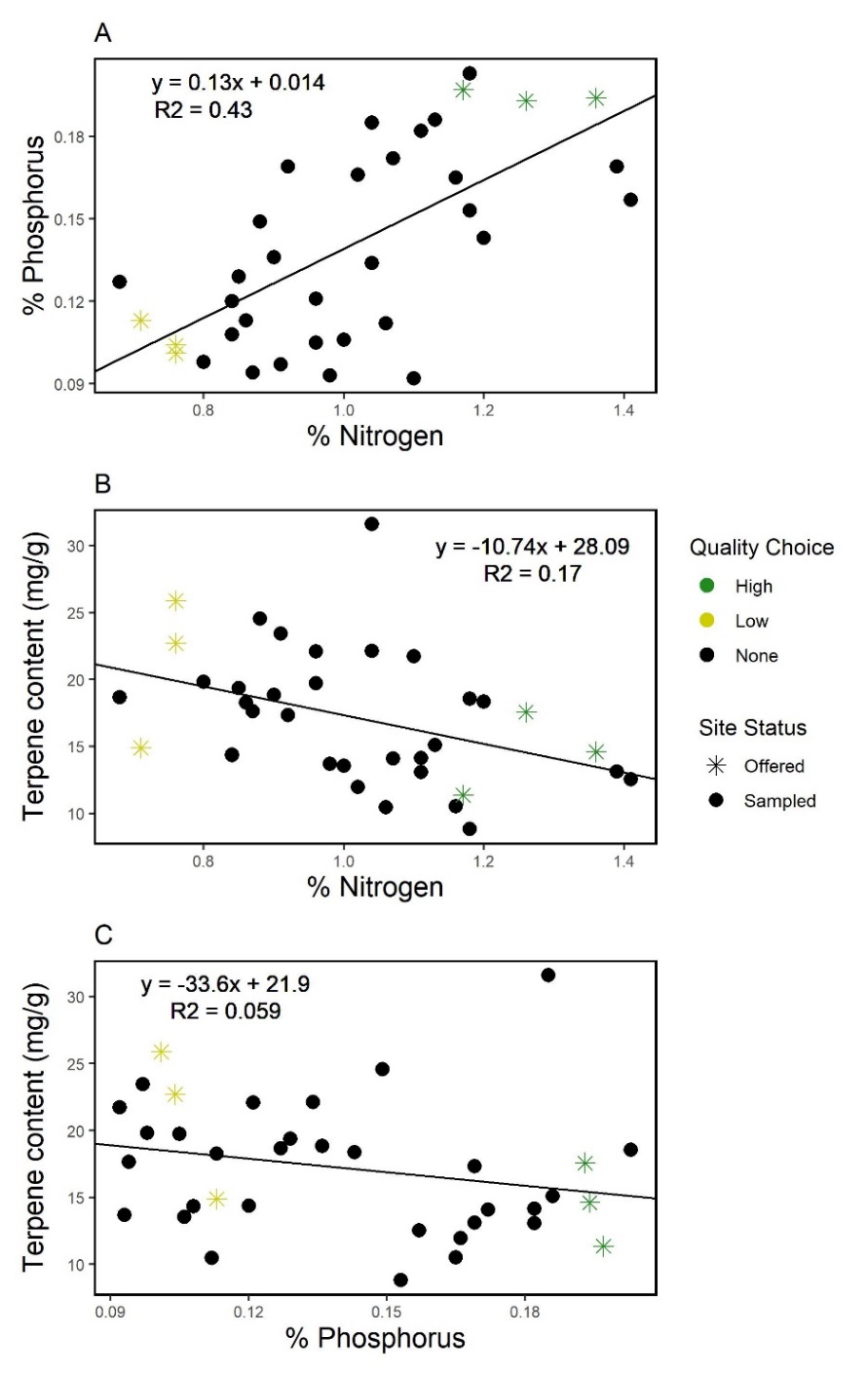

Figure A2. Relationships between black spruce A) nitrogen and phosphorus compositions, B) nitrogen compositions and plant secondary compound concentrations (PSC), and C) phosphorus compositions and PSC concentrations from all trapping grid sites sampled in 2017 (n = 36). Stars represent sites where we clipped branches for cafeteria experiments during the autumns of 2018 and 2019, either high-quality (green) or low-quality (yellow) black spruce.

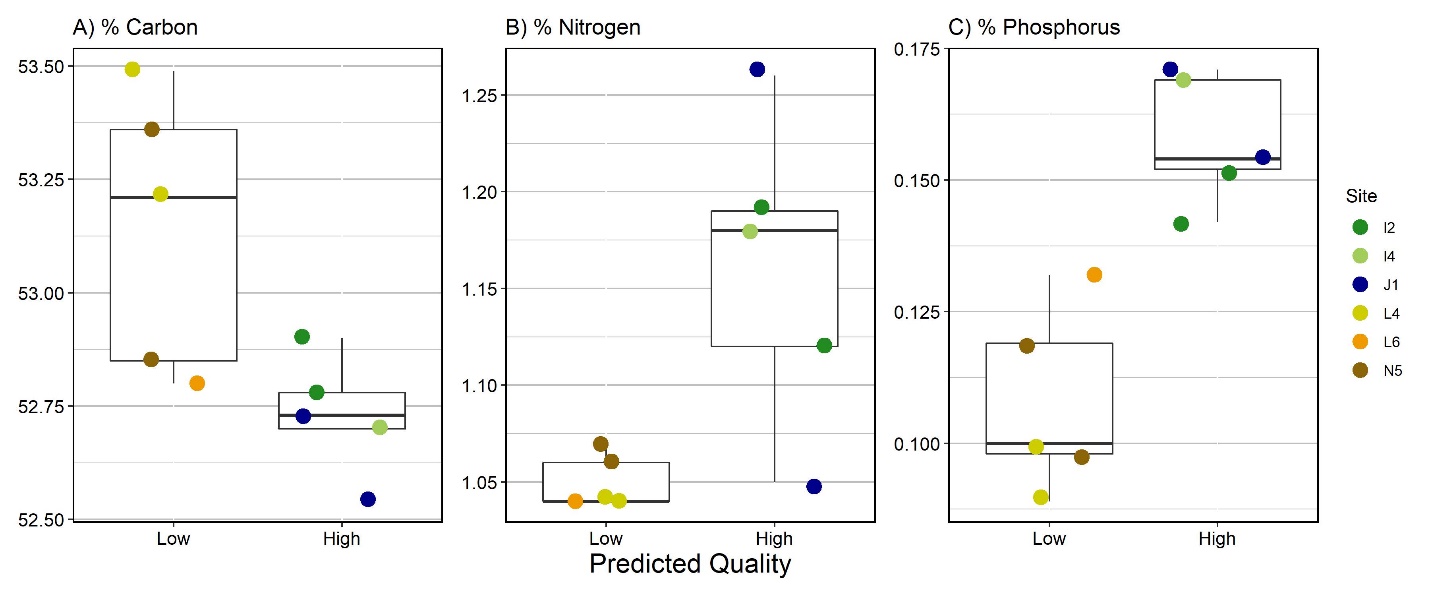

Figure A3. Carbon (A), Nitrogen (B), and Phosphorus (C) compositions of black spruce (*Picea Mariana*) subsamples (n = 10) from offerings for snowshoe hare cafeteria experiments in 2019 compared by their predicted nutritional rank and coloured by sample site. Based on sampling from the summer of 2016, we predicted that spruce subsamples from high-quality areas would have higher N and P compositions and lower C compositions than low-quality areas.

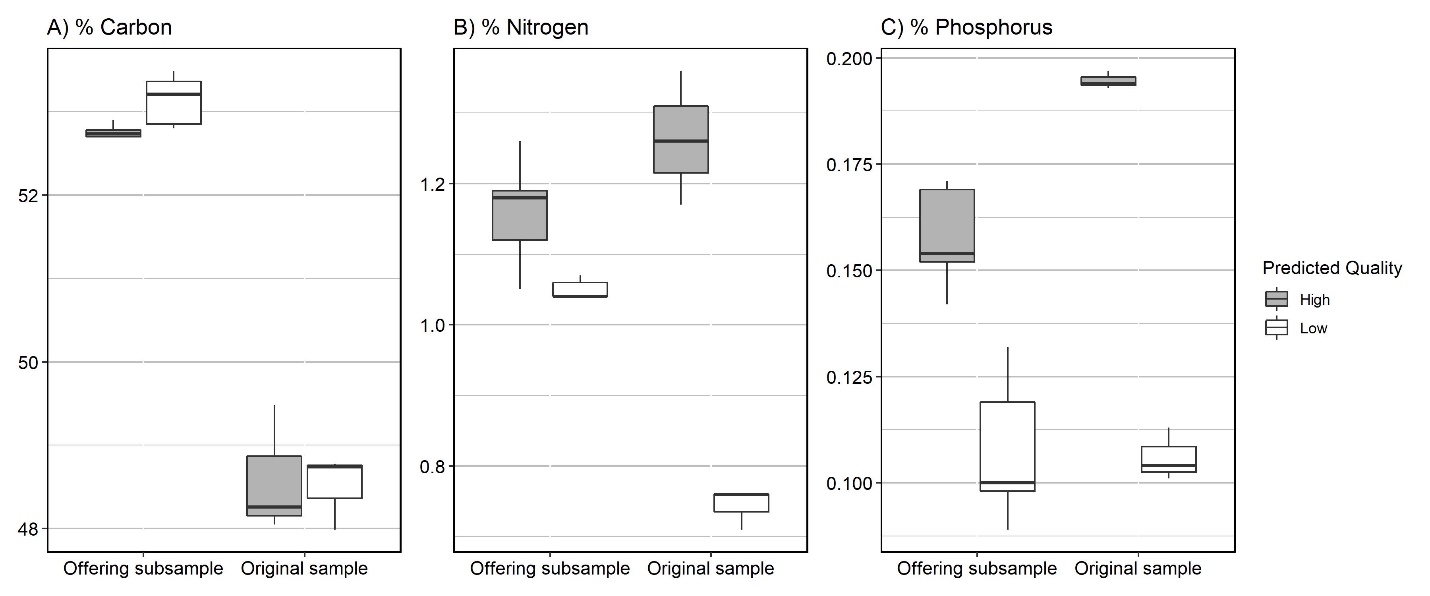

Figure A4. Carbon (A), Nitrogen (B), and Phosphorus (C) compositions of black spruce (*Picea Mariana*) subsamples (n = 10) from offerings for snowshoe hare cafeteria experiments (fall 2019) compared to original samples (summer 2016) from the same set of sample sites, either high-quality areas (high N and P; shaded) or low-quality areas (low N and P; not shaded). We predicted that cafeteria offering subsamples would show a similar trend to the original samples. There is a higher concentration of C in offered samples compared to original samples and we assume this to be due to an increase in indigestible carbohydrates due to phenology and lignification.

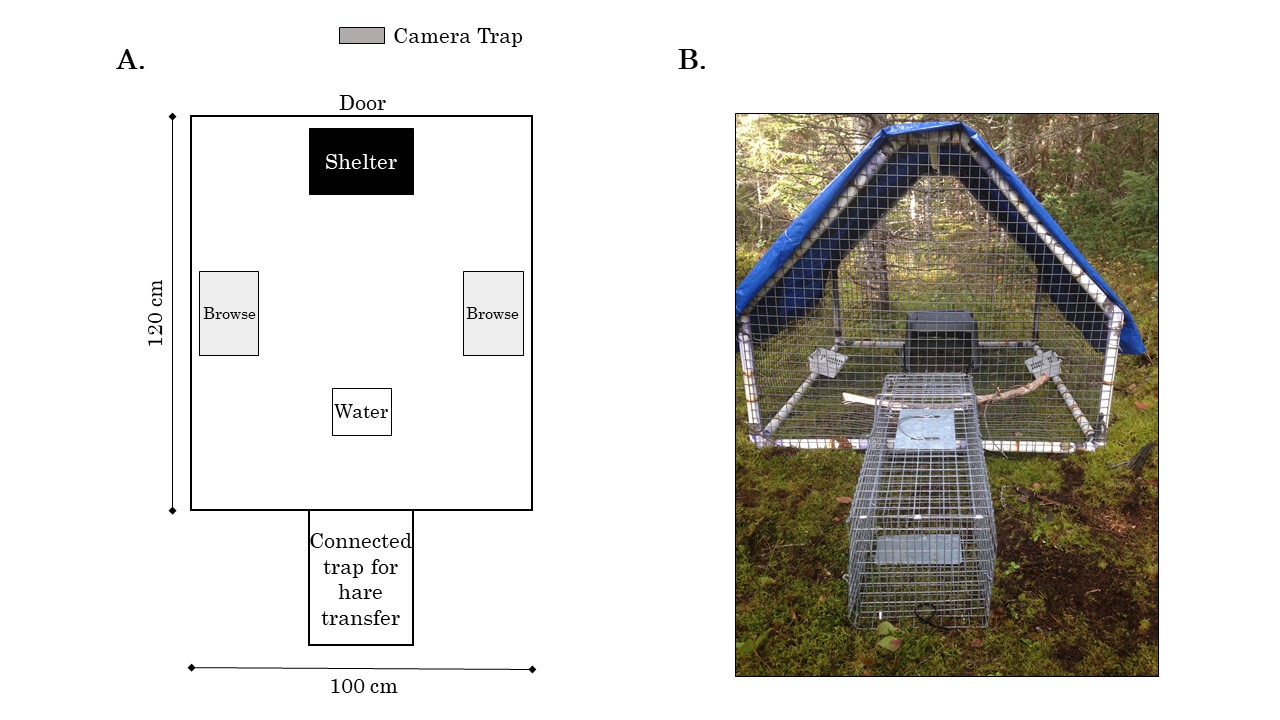

Figure A5. Overhead diagram (A) and photograph (B) of a snowshoe hare cafeteria enclosure.

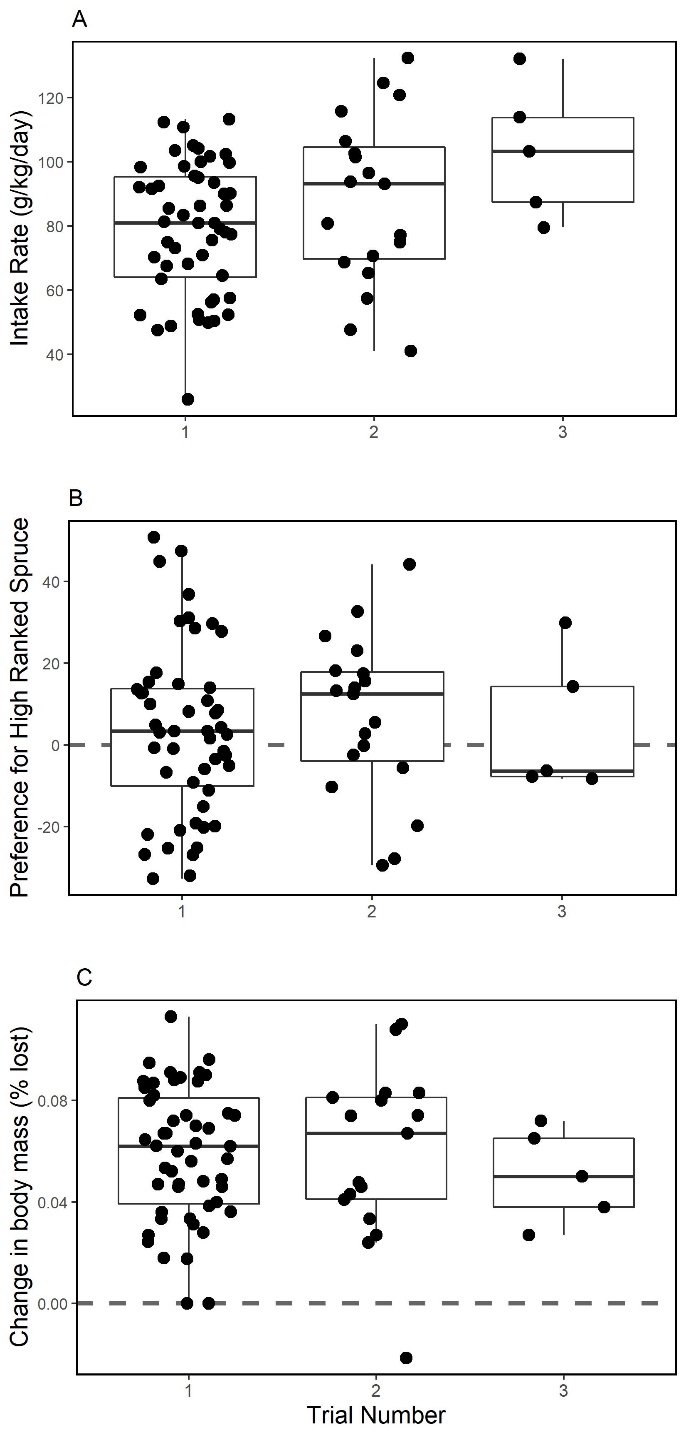

Figure A6. A) The intake rate (IR; g·kg^-1^·day^-1^; n = 75) of all offered spruce (high and low quality), B) preference for higher ranked spruce (IR high ranked – IR low ranked; n = 75), and C) weight loss (% lost per day; n = 73) by individual snowshoe hares during 24-hour captive cafeteria experiments in 2018 and 2019 compared to the trial number. The dashed zero lines in panels B and C represents no preference between spruce ranks and no weight loss respectively.

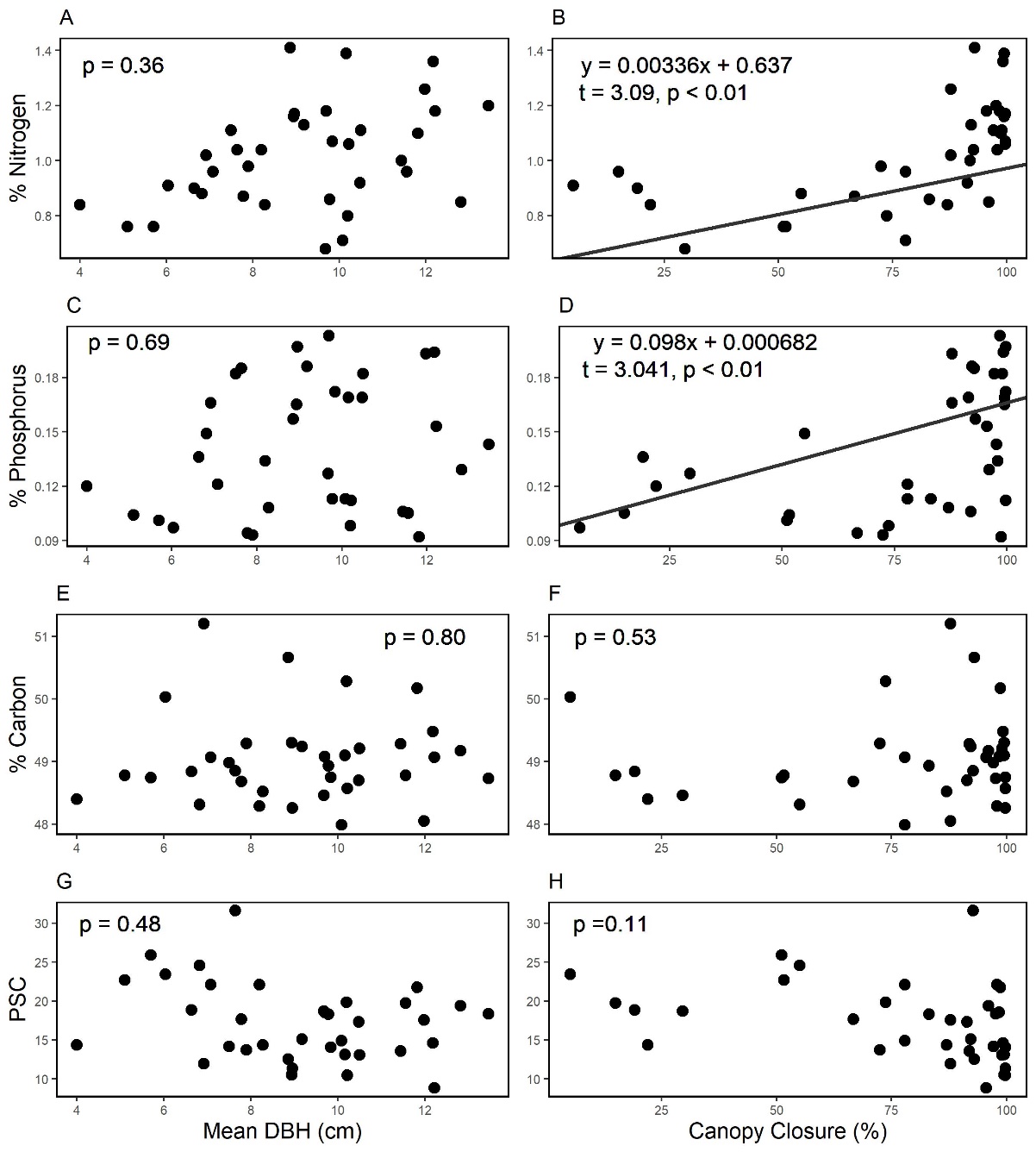

Figure A7. Black spruce nitrogen compositions (A and B), phosphorus compositions (C and D), carbon compositions (E and F), and plant secondary compound (PSC) concentrations (G and H) sampled at 36 sites across the trapping grid plotted against site mean tree Diameter Breast Height (DBH; A, C, E, and G) and canopy closure (B, C, F, and H). Linear regressions are shown for significant effects (p < 0.05). Collectively, DBH and canopy closure explained 36.0%, 25.1%, and 26.5% of spruce nitrogen, phosphorus, and PSC variation, respectively.

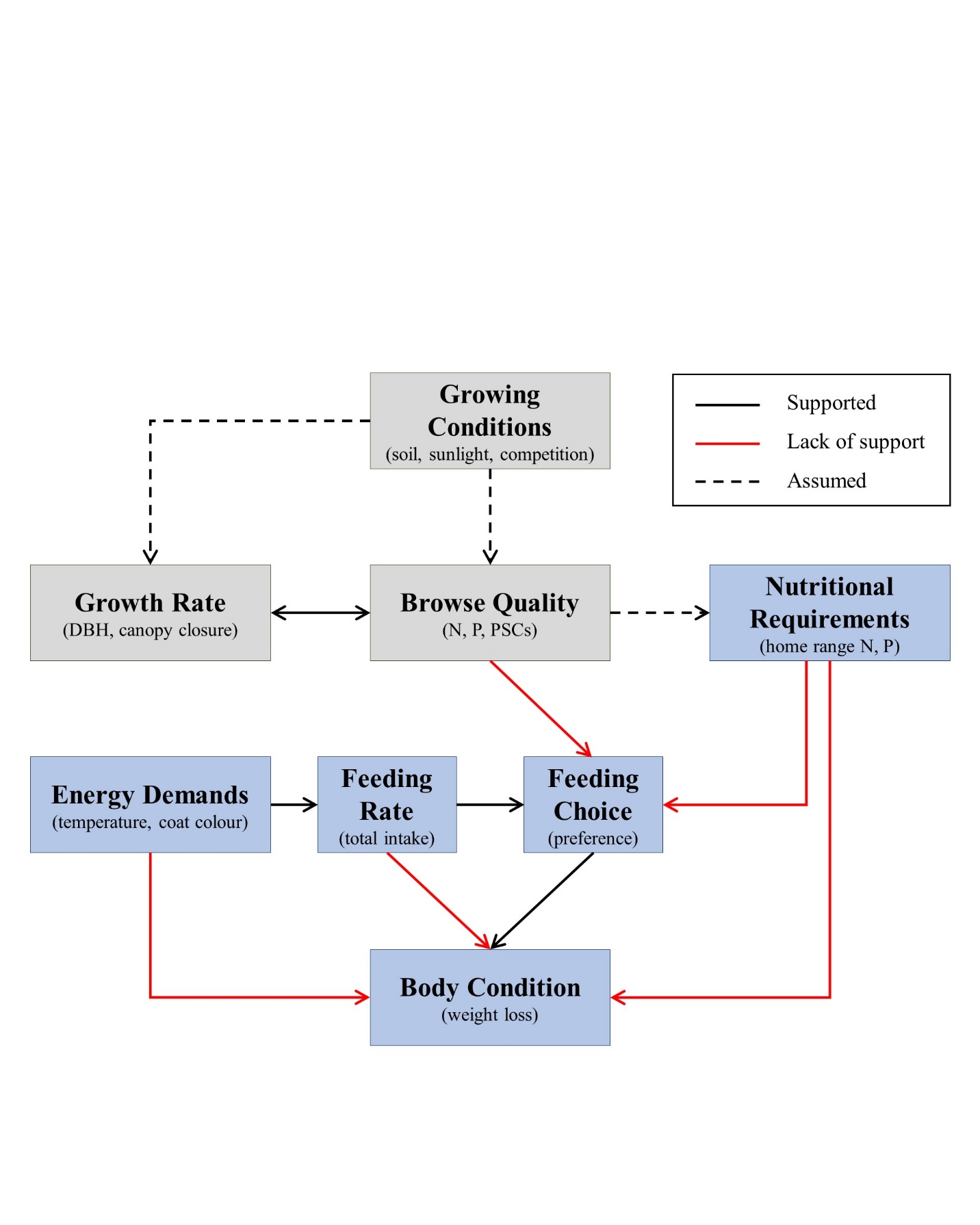

Figure A8. All predictions linking plant (grey) growing conditions and quality to herbivore (blue) feeding and body condition responses of our study using black spruce (*Picea mariana*) and snowshoe hare (*Lepus americanus*). Predictions are coded as either supported (solid black), not supported (solid red), or assumed but not tested in this study (dashed black).

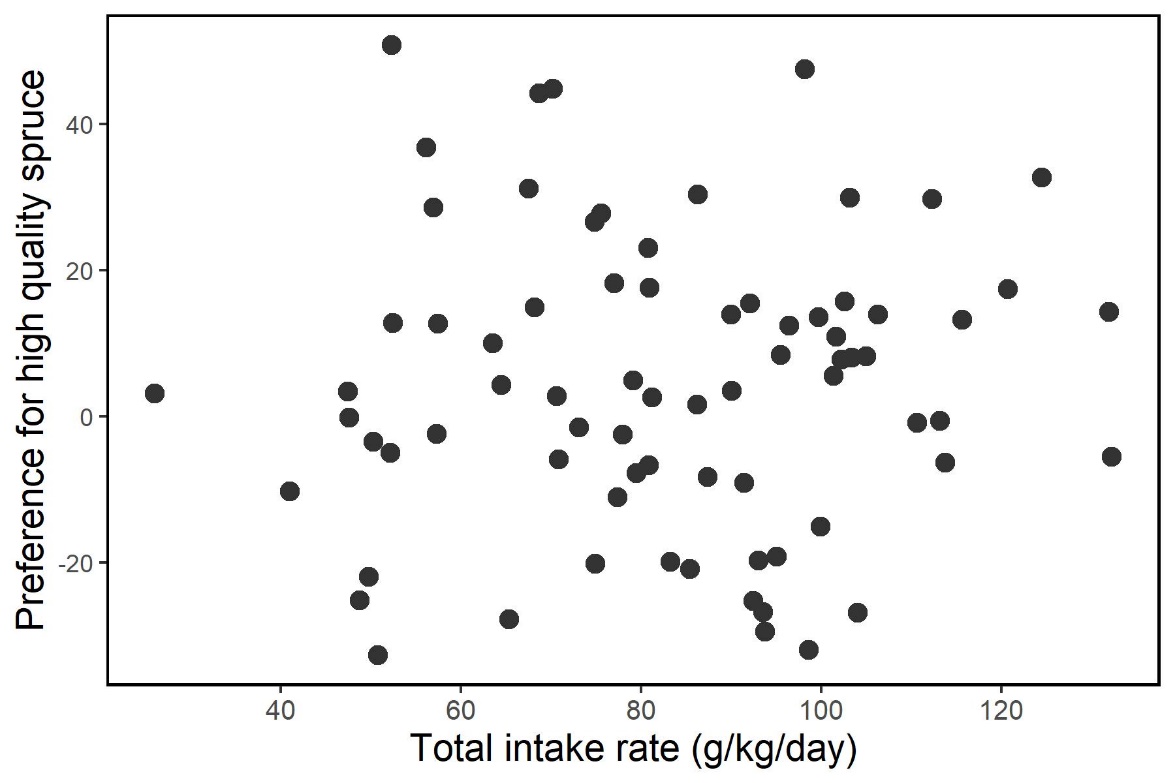

Figure A9. Preference for high-quality spruce (high-quality intake rate – low quality intake rate; g·kg^-1^·day^-1^) by snowshoe hares during cafeteria experiments, which tested their feeding response to two qualities of black spruce, in relation to their total spruce intake during experiments. The relationship between intake rate and preference (preference = 1.13 + 0.038*intake rate) proves to not be significant (t = -1.34; p = 0.18). This relationship was not included in our AICc models.
